# QuasiFlow: a bioinformatic tool for genetic variability analysis from next generation sequencing data

**DOI:** 10.1101/2022.04.05.487169

**Authors:** Pedro Seoane, Luis Díaz-Martínez, Enrique Viguera, M. Gonzalo Claros, Ana Grande-Pérez

## Abstract

Populations of RNA and ssDNA viruses within their hosts contain a heterogeneous collection of variant genomes known as quasispecies. Large variability in mitochondrial DNA has also been found within the same organism, drawing an interesting parallel between the two situations. The advent of next-generation sequencing technologies facilitated studying genetic variation, but many open-source bioinformatic tools have to be combined in a non-trivial approach. Here it is presented QuasiFlow, a workflow based on well-stablished software that extracts reliable mutations and recombinations, even at low frequencies (~10^-4^), provided that at least 250 million nucleotides are analysed. Accurate prediction of mutations and recombinations has been demonstrated with synthetic reads and with in vitro rolling-circle amplification of a plant geminivirus. An in-depth analysis of viral quasispecies was performed and QuasiFlow revealed the coexistence in the plant of three virus genomes and distinct recombinations between some of them. Human mitochondrial variants were also investigated and high level of heteroplasmy (75%) was confirmed, and the relation between low-frequency heteroplasmy (0.1- 0.2%) and some human diseases, regardless of sex, was established. Hence, we propose that QuasiFlow may find use with known and emerging viruses to reveal evolutionary jumps and co-infections, with mitochondrial DNA to detect relevant heteroplasmy would otherwise be elusive, or even in other population studies such as those considering single cell sequencing.

## INTRODUCTION

RNA and ssDNA viruses, including SARS-CoV-2, exist as highly variable and heterogeneous collections of mutated and recombinant genomes known as virus quasispecies (Lauring and Andino 2010; Domingo et al. 2021). The term ‘quasispecies’ means that a viral genome is indeed a weighted average of a large number of different but related individual genotypes, called haplotypes (Domingo and Perales 2019; Domingo et al. 2021). Haplotypes differ not only in point mutations, since it is well known that many viruses can recombine resulting in the shuffling of beneficious and deleterious variations that can be separated and tested as individual units in new combinations. High variability has also been found in mitochondrial DNA (mtDNA) that has been related to human mtDNA disorders that affect one of 4300 individuals (Gorman et al. 2015). The frequency in which the mtDNA variants (point mutations and indels) are found determines that they are considered heteroplasmy, if different variants of the mtDNA coexist in the same cell, tissue, organ or organism, or homoplasmy, in case all mtDNA copies share the mutation. Heteroplasmy and homoplasmy have been related to disease and ageing in humans (Stewart and Chinnery 2015).

Studies on virus quasiespecies usually required laborious and expensive laboratory efforts. Fortunately, the advent of next-generation sequencing (NGS) technologies enabled the fast and cheap detection of any minor variant within a huge population of genomes (Bull et al. 2016; Posada-Cespedes et al. 2017) and discern all single nucleotide variations (SNVs) in any quasispecies to define a haplotype (viral variant) without the need of culture or cloning prior to sequencing (Zukurov et al. 2016). However, analysing NGS data can be an awkward endeavour starting from the available open-source, cutting-edge software with incompatible formats to study recombination, variant calling and haplotypes (Cacciabue et al. 2020). Additionally, quasispecies studies must deal with sequencing errors, whose frequency lays close to the viral replication error rates (10^-3^-10^-5^ errors per nucleotide).

To fill this gap, here we describe QuasiFlow, a bioinformatic tool based on well stablished software to study genetic variability from NGS data that is able to distinguishing real variations from sequencing errors. Accurate and precise detection of low-frequency mutations, genetic variability and recombination is achieved in viral quasispecies and other highly variable genomes like mtDNA. Its companion QuasiComparer facilitates the interpreation of multiple analyses on comparable samples. Both pipelines together can extract the complete genetic variability provided that more than 250 million nucleotides were sequenced.

## RESULTS

### Stringent filtering in QuasiFlow provides a reliable detection of low-frequency genetic variations

The performance of QuasiFlow workflow (Fig. 1) was investigated using four datasets (mut_rec, mut_REC, MUT_rec and MUT_REC, where ‘MUT/mut’ means mutations, ‘REC/rec’ means recombination, lower case indicates absence, and upper case indicates presence) of synthetic reads mimicking an Illumina sequencing project (see Materials and Methods for details). Read rejection rate was 0.72 in all four groups (Fig. 2A, RejectedReads) due to the strict requeriments in the read-preprocessing step to avoid the confusion of sequencing errors with mutation events. Consequently, the useful reads (Fig. 2A, ReferenceReads) presented a recovering index of 0.27 for recombinant free datasets (mut_rec and MUT_rec) with a slight decrease to ~0.25 where only recombinations (mut_REC) or recombinations together with mutations (MUT_REC) were present, respectively. Mapping rate of useful reads was nearly 1.00 for the samples without mutations (Fig 2A, MappedReads) with a slight decrease to 0.98 when reads contain artificial mutations, indicating that the accumulation of sequence changes causes the alignment rejection by the mapper with highly strict parameters. Concerning recombination, untangled datasets (mut_rec and MUT_rec) showed an expected 0.0 rate for true and false recombination events (Fig 2A, TrueRECs and FalseRECs) while most (0.86) recombination events were correctly predicted in both mut_REC and MUT_REC. The striking value of false recombinants in the MUT_rec sample is due to one single unexpected recombination event. The few cases of false recombinations detected in MUT_REC (Fig. 2A, FalseRECs) were manually inspected revealing that they were true recombinants produced 4-5 nt apart from the expected nucleotide, indicating that stringent validations of QuasiFlow detects true recombinations, even if they are not at the expected position and should not affect when real-world data are used. Concerning mutations, it results that unmutated datasets (mut_rec and mut_REC) contained no mutation per read (Fig. 2A, TrueMUTs), while mutated datasets (MUT_rec and MUT_REC) were 0.863 and 0.831, respectively. Taken together, Fig. 2A demonstrated that QuasiFlow can predict very accurately the mutation or recombination frequency in a population using only the subset of useful reads.

**Figure 1:**
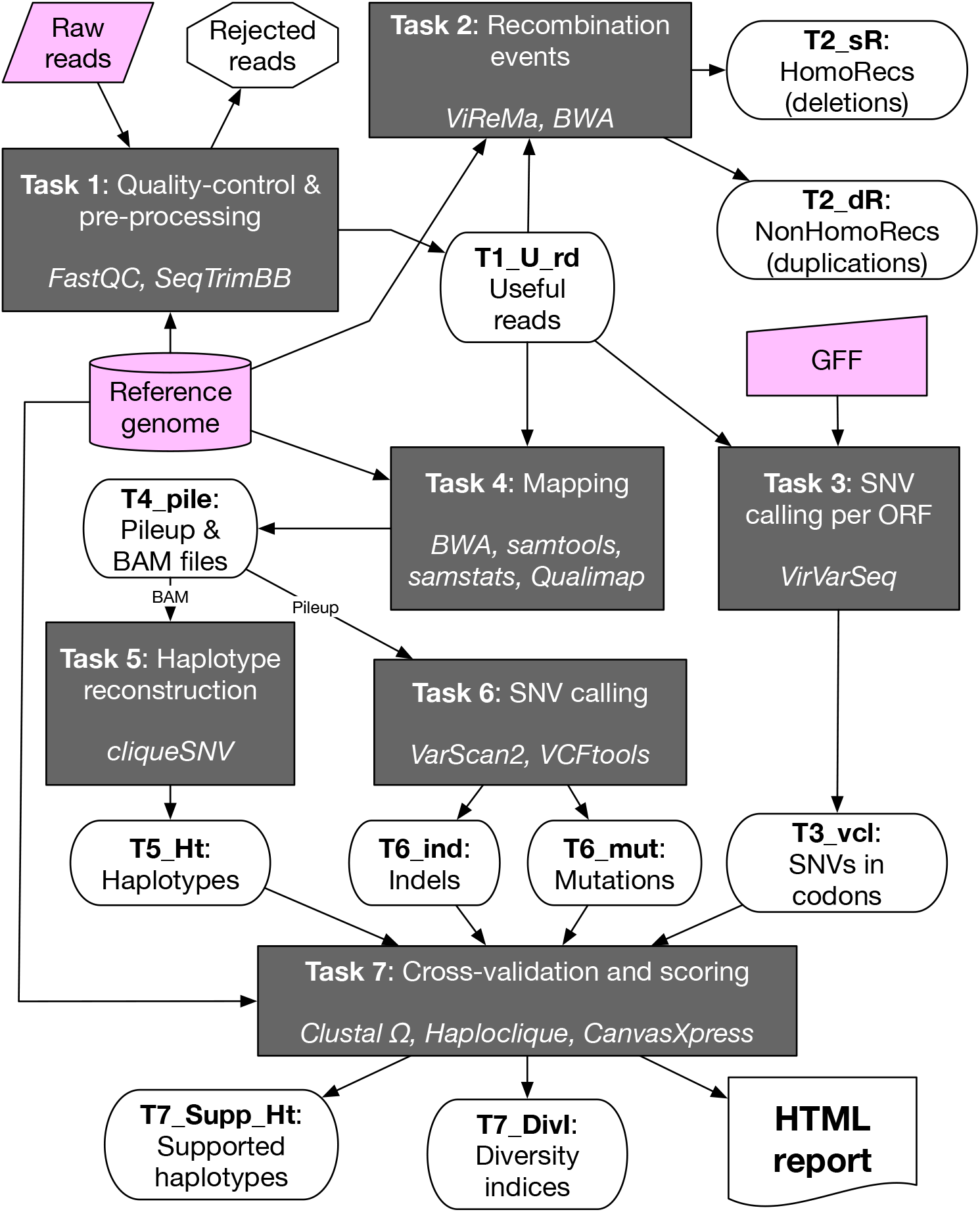
Conceptual representation of the QuasiFlow workflow for quantifying genetic variability using high-throughput sequencing reads. The seven tasks, described in detail in Materials and Methods, are grey rectangles, where the relevant bioinformatic tools are indicated in italics. Magenta boxes refer to QuasiFlow input files. White rounded boxes correspond to input/output data recoverable when QuasiFlow finishes; their title containing the task number will facilitate its citation throughout the text. Solid arrows indicate information flow.

**Figure 2:**
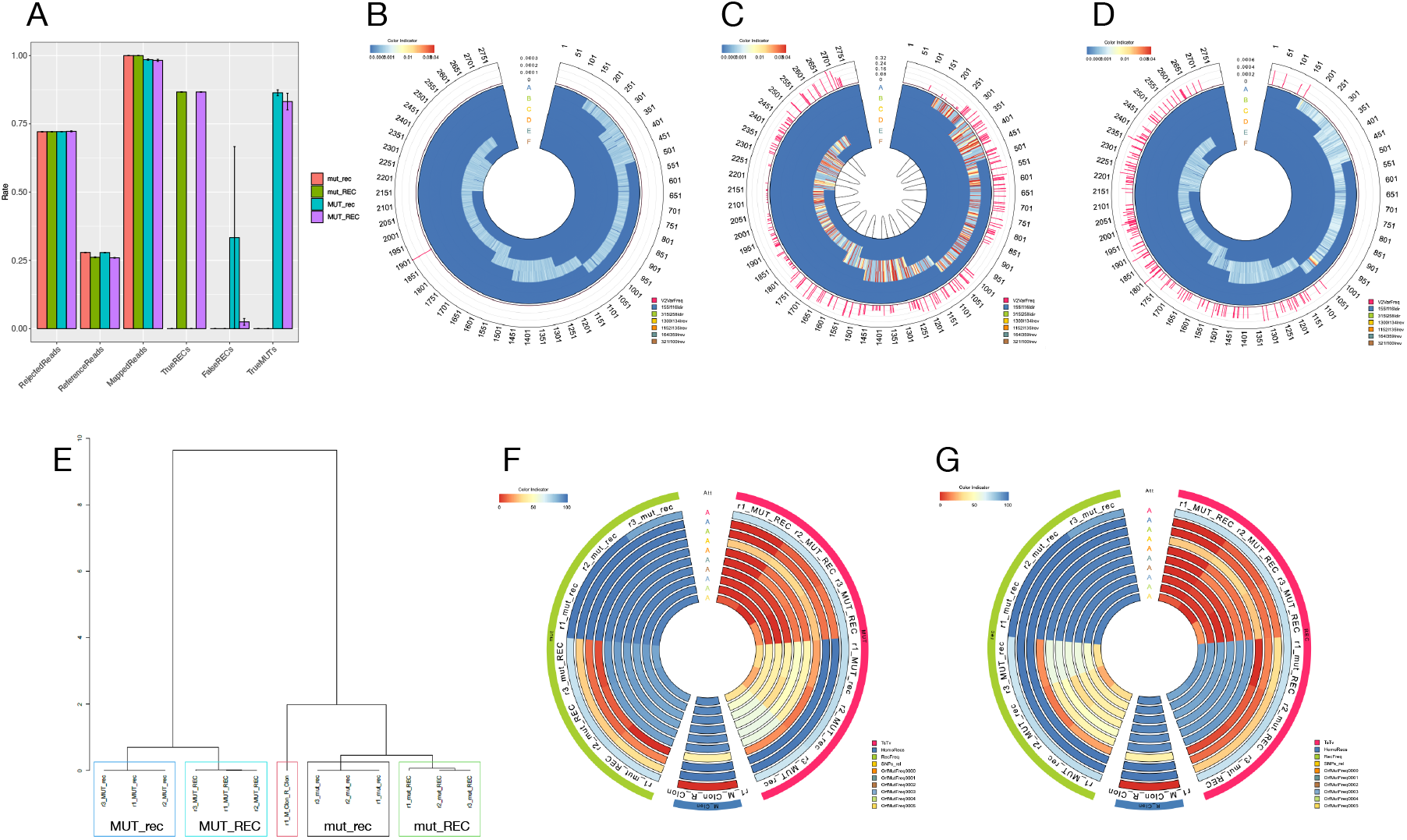
Quasiflow and QuasiComparer performance. (**A**) Benchmarking QuasiFlow with synthetic reads. Bars represent means of three replicates with their standard error. Colours are indicated in the legend and correspond to the four synthetic datasets. “RejectedReads” are reads removed due to quality, complexity or length issues. “ReferenceReads” are reads devoid of artefacts (useful reads) and therefore suitable for genetic variance analyses. “MappedReads” are useful reads mapped onto reference sequence of TYLCV-Mld. “TrueRECs” represent recombination events that were predicted at the expected position. “False RECs” gather those recombination events that were not detected at the expected position. “True MUTs” correspond to mutation events predicted at the expected position. “Haplotypes” are those retained after task 7 cross-validation (T1_Supp_Ht in Fig. 1). Variation profile of replicates from synthetic mut_rec (**B**) and MUT_REC (**C**), and RCA-amplified TYLCV-Mld clone (**D**), where the outer track (V2VarFreq) corresponds to the frequency of variants called by Varscan2, the other six inner tracks are VirVarSeq mutation frequencies determined in the six TYLCV-Mld frames), and the internal links represent origin and target positions of the recombinations identified by ViReMa. (**E**) Hierarchical clustering after PCA of three replicates of each synthetic dataset and the TYLCV-Mld, where the different replicates are clustered together and surrounded by color boxes, and the single TYLCV-Mld sample is boxed in red. Analysis of variance (ANOVA) for the mutation (**F**) and recombination (**G**) rates in the same samples of panel E. The outer ring groups samples by factor: in green are synthetic samples that lack of inserted mutations (mut) or recombinations (rec), in pink are gathered all samples with inserted mutations (MUT) or recombinations (REC), and in blue is the TYLCV-Mld infectious clone sample. Inner rings represent the most significant variables, where the colour spectrum represents the relative value of the variable in the range of 0 (blue) to 100% (red), as indicated in the colour indicator. The QuasiFlow HTML report for mut_rec (*1-mut_rec.html*) and MUT_REC (*2-MUT_REC.html*) synthetic reads, the QuasiComparer complete comparative report for all synthetic reads (*3-Comparative_Report_SyntheticReads.html*), as well as for the control of TYCLV-Mld replicated in vitro (*4-Control_TYLCV.html*) can be inspected in Supplementary File 1.

The whole variation profile was then investigated (Fig. 2B-D). As expected, the control group of mut_rec presented no mutations and no recombination events, being consistent with a very low frequency of genetic variability (Fig. 2B). For high variation profiles such as MUT_REC (Fig 2C), 261 new mutations with frequencies ranging from 0.31 to 1.72 × 10^-4^, and 13 recombination events were detected. This illustrates that QuasiFlow can find both low- and high-frequent mutation and recombination events. A test using a real-world dataset from a RCA-amplified infectious clone of the begomovirus TYLCV-Mld (tomato yellow leaf curl virus, mild strain), where low mutation frequency is expected as well as absence of recombination, 72 mutations with frequencies ranging from 4.2 × 10^-4^ to 2.2 × 10^-4^ were detected and no recombination events were identified (Fig. 2D). The apparent hotspot detected from the nucleotide 1110 to 1128 for the ORF so called 315-258-dir (Fig 2D, track B) is due to the quasi-dimer (1,7-mer) nature of infectious clone compared to the reference genome, being the region where digestion and fusion was produced and is not identical in all virus haplotypes. Since the RCA-amplified TYLCV-Mld (Fig 2D) presents values closer to mut_rec (Fig 2B) than to MUT_REC (Fig 2C), it can be suggested QuasiFlow was able to detect the low amount of variation introduced by bacterial cloning and RCA manipulation, and that this low level of variability would have a negligible impact on genetic variation. This also supports that mutation and recombination events and the resulting haplotypes provided by QuasiFlow should be considered reliable, derived from real biological events and not from experimental artefacts.

Quasispecies studies usually require comparison of several samples. This has been facilitated by another workflow called QuasiComparer. When samples mut_rec, mut_REC, MUT_rec, MUT_REC and TYLCV-Mld where compared (*3-Comparative_Report_SyntheticReads.html* in Supplementary File 1), most (67.54%) of the total variance in PCA seems to depend on mutations, which explains why the TYLCV-Mld sample clusters with mut_rec and mut_REC datasets (Fig. 2E), in agreement with resultis in Figs. 2B and 2D. The same behaviour can be inferred after ANOVA taking into account both mutations (Fig. 2F) or recombinations (Fig. 2E) as factors, the same 4 variables being highly significant for both factors (*P* < 0.005). Since the TYLCV-Mld sample shows again a pattern close to datasets without recombination events and without mutations, it is suggested that QuasiComparer easily illustrates the classifation of known samples as expected, and serves to group samples based on their biological origin, being a great help to interpret results.

### Quasiespecies reconstruction of original and recombinant geminiviruses in a tomato plant

QuasiFlow performance detecting recombinant viruses was illustrated using geminivirus. MiSeq data from a rolling circle amplification (RCA) product of a TYLCV-Mld agroinfecting *Solanum lycopersicum* plant was analysed. Viral genetic variation was evaluated with the incidence, abundance and functional diversity indexes described in methods presented in Table S1 of Supplementary File 2. These values are characteristic of highly diverse viral populations (Gregori et al. 2016), since (i) the mutation frequency was 1.008 × 10^-2^ mut/nt, two orders of magnitude higher than that of other geminiviruses (~10^-4^ mut/nt) (Isnard et al. 1998; Sanz et al. 2000; Ge et al. 2007; Duffy and Holmes 2008); and (ii) the Shannon index reached 0.6. The consensus sequence differed by 32 SNVs from the reference sequence (TYLCV-Mld) and its mutation frequency per nucleotide position clustered into two well differentiated regions (Figure 3A). The first, from positions 1 to 1000 and includes 21 changes with a mutation frequency of 0.85 ± 0.03. A second region, from position 1201 to 2791, included changes with mutation frequencies of 0.95 ± 0.03 and 0.69 ± 0.02. Between the two regions, at positions 1000-1200 a hostpot was found with a mutation frequency of 0.45. A total of 31 haplotypes (Hp) were reconstructed (Figure 3B), including one (Hp1) identical to the reference sequence. Hp12, Hp27 and Hp31 accounted for 80% of the total. Hp12 showed an identity of 98% with TYLCV-Mld and represented 40% of the total. Interestingly, blast analysis indicated that Hp12 was identical to Almería strain of TYLCV (TYLCV-Almería, AJ489258.1) from nt 1-1054 whereas from nt 1055-2791 was identical to Malaga strain (TYLCMaV, AF271234.1). The recombinant nature of Hp12 was confirmed by RDP analysis (Fig. 3C) and suggested a triple infection within the agro-infected plant. *In silico* results were confirmed by restriction analysis (Fig. 3D) of the RCA product. *Kpnl* releases the monomer of TYLCMaV and *ClaI* cuts TYLCV-Almería into two fragments and linearises TYLCV-Mld. This assay revealed the existence of the three viruses in the tomato plant. In addition, *bona fide* recombinant genomes between TYLCV-Almería and TYLCMaV (Fig. 3E) were amplified with specific primers. Finally, since haplotype reconstruction showed a high frequency of genomes carrying changes that belonged to the TYLCMaV sequence, the relative abundance of each virus was investigated. The analysis of the coverage of each parental virus (Figure 3F), using an artificial reference sequence with the genome of the three viruses, showed that most of the variants belonged to TYLCMaV, followed by TYLCV-Almería, while TYLCV-Mld was almost undetectable, suggesting a fitness advantage of TYLCMaV.

**Figure 3.**
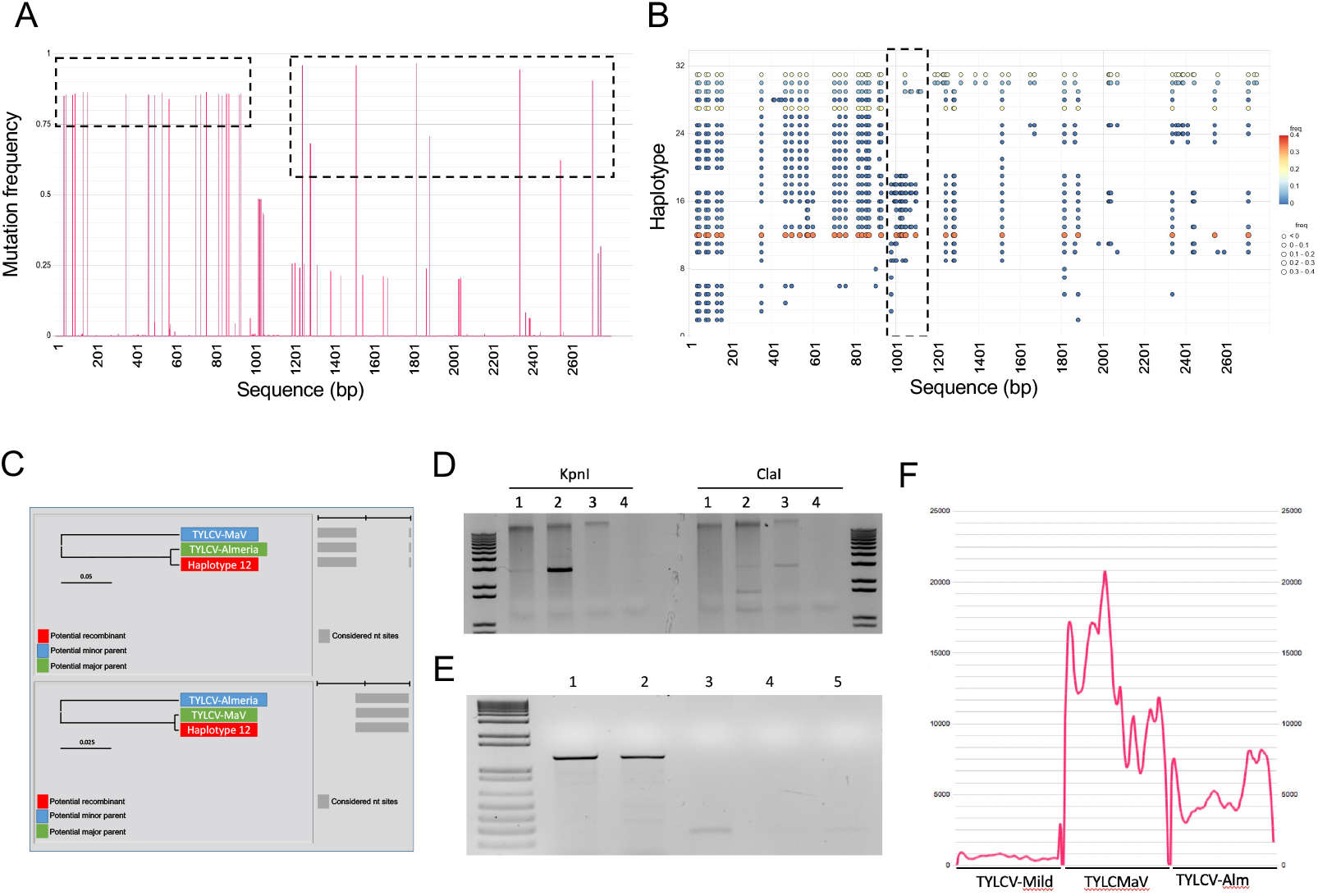
Variation analysis of a TYLCV-Mld population after infection of tomato plants. **(A)** Nucleotide mutation frequency profile, where black dashed lines indicate regions that group SNPs of similar frequency, one containing mutations with two different frequencies. Only mutations with a frequency over 0.5 were considered for the consensus. (**B)** Haplotype reconstruction of the TYLCV population. Circles depict mutated positions on each haplotype respect to the reference sequence TYLCV-Mld. Coloring indicates haplotype frequency. Of the 31 different reconstructed haplotypes, haplotype 1 (Hp1) is identical to the consensus sequence. The dashed squares frame regions with high variability among haplotypes. (**C)** RDP4 analysis showing Haplotype 12 as a recombinant with the first part of its sequence (gray rectangles, upper panel) of TYLCV-Almería and the second half of TYLCMaV (lower panel). **(D)** Restriction analysis of the RCA product from the tomato sample: 1/100 dilution (lanes 1) and a 1/10 dilution (lanes 2); positive control, TYLCV-Mld monoinfected tomato (lanes 3); negative control (lanes 4). (**E)** Identification of a *bona-fide* recombinant between TYLCMaV and TYLCV-Almería in the TYLCV-Mld infected tomato by PCR. A region of 1.4 kb of the recombinant was amplified in a 1/100 dilution (lane1), undiluted DNA (lane 2); positive controls: tomato monoinfected with TYLCV-Mld (lane 3), TYLCMaV (lane 4) and TYLCV-Almería-(lane 5). (**F)** Nucleotide coverage of the infected tomato sample using a TYLCV-Mld, TYLCMaV and TYLCV-Almería chimaera as a reference sequence. The QuasiComparer complete comparative report for tomato infected plants with TYCLV (*5-Tomato_TYLCV-invected.html*) can be inspected in Supplementary File 1.

### Study of mitochondrial variants to establish a relationship between low-frequency heteroplasmies and human diseases

Within eucariotic cells, a population of mitochondrial genomes exists, each genome harboring variations called variants of mitochondrial DNA (mtDNA). We analysed MiSeq data of 47 human mtDNA samples from blood, lymphoblastoid cell lines and saliva. QuasiFlow detected 10 400 variants of which 14.5% (1 508 variants) were homoplasmies and 85.5% (8 892 variants) corresponded to heteroplasmies. Nucleotide changes (Supplementary File 2, Table S2) were biased towards transitions A → G and T → C in all samples, followed by C → T and G → A. No transversions were found in 12 of the 47 samples. Looking at heteroplasmies, a remarkable 80% had very low frequency, that is, between 0.1% and 0.2%, and the rest (20%) had a frequency no higher than 2% (Fig 4A). These results suggest that mtDNA populations in these samples maintain a very low level (below 0.3%) of heteroplasmies while homoplasmies were very frequent (greater than 98%) with a stricking absence of intermediate cases. When the relationship between mtDNA variants and disease was inspected, 451 variants were associated with different diseases (Supplementary File 2, Table S3). Interestingly, more than 75% of these mtDNA variants were heteroplasmies.

**Figure 4.**
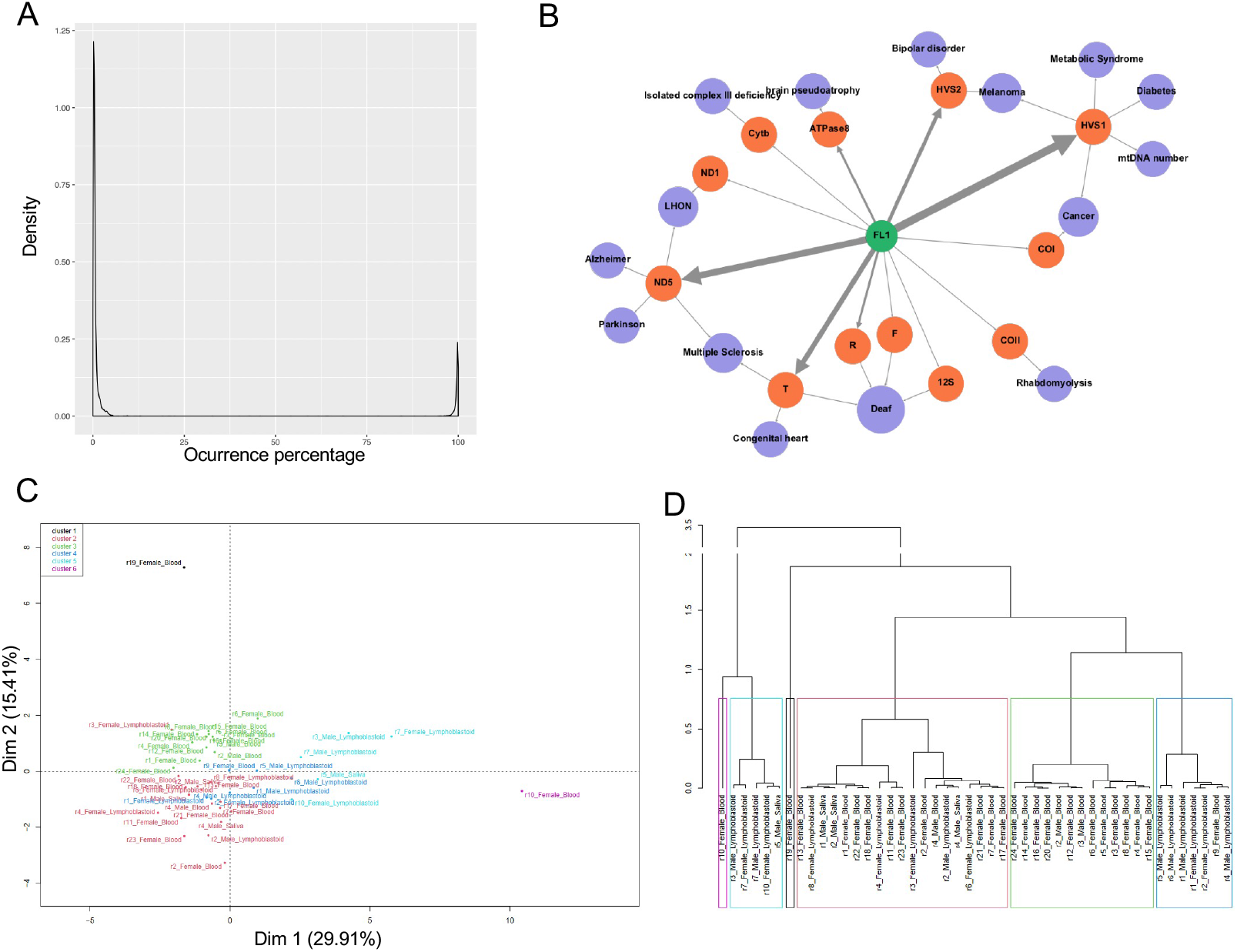
Variation analysis of mtDNA populations from different human tissues. **(A)** Average distribution of mtDNA variants in the 47 samples analyzed showing very low frequency of heteroplasmies (below 0.2%) and homoplasmies, but without intermediate frequency variants. **(B)** Mutation network of sample 1 of the lymphoblastoid cell line (FL1, green node). ORFs with heteroplasmies (orange nodules) are linked to diseases (purple nodules) that have been associated with these mutations. The thickness of the line is proportional to the frequency of the heteroplasmy. (**C)** Sorting the different mtDNA samples by PCA. The colors correspond to the different clusters formed from the analysis of the first four dimensions of the PCA. (**D)** Cladogram showing the five clusters from the PCA of panel D. Blood samples group in the red and black clusters, linfoblastoid samples in the blue and red clusters, and saliva samples are dispersed throughout the three main clusters. The QuasiComparer complete comparative report for the mtDNAs (*6-Comparative_Report_Mitochondria.html*) can be inspected in Supplementary File 1.

Using the data retrieved from MITOMAP a network was constructed for each sample illustrating the relationship between mtDNA variants per ORF and disease. An example is shown in Figure 4B with biological replicate 1 of a female lymphoblastoid sample. Note that 26 homoplasmies were related to brain syndromes such as Alzheimer’s, Parkinson’s, deafness, bipolar disorder, multiple sclerosis, Leber hereditary optic neuropathy (LHON) or cerebral pseudoatrophy. Remakably, 310 heteroplasmies, of which 300 had a frequency below 1%, were related to the same brain syndromes. These findings highlight the importance of detecting low frequency mtDNA heteroplasmies for disease association.

The impact of sex and tissue factors were inspected using QuasiComparer. No significant differences (P < 0.05; Table S4 in Supplementary File 2) were detected among the 47 samples for any of the factors, but differences become evident by PCA for tissue factor (Fig. 4C), that explained 48% of the total variance. In fact, hierarchical clustering grouped samples in four clusters (Fig. 4D) two of them by tissue, while no association between sex and mtDNA variants was found. The blue- and cyan-boxed clusters in Fig. 4D contained mainly lymphoblastoid samples and the red-boxed cluster was enriched with blood samples. Differences between these clusters were found significant (*t*-Student, *P* < 0.05).

### Genetic variability requires sequencing of 250 million nucleotides in TYLCV

To present a robust and accurate assessment of mutation and recombination frequencies, the QuasiFlow/QuasiComparer analysis must rely on the whole genetic variability, and this is clearly dependent on the number of reads for the low frequent SNVs. Hence, the number of mutations detected using MiSeq or HiSeq platforms on the same sample was compared depending on the total number of nucleotides sequenced. A sigmoid behavior was observed using both platforms (Figure S1 in Supplementary File 2), reaching a plateau (that is, mutations at all frequencies were detected) when nearly 250 million nucleotides were sequenced. It is reassuring that both sequencing platforms require the same number of nucleotides sequenced to saturate the analysis, corresponding to ~1.5 millions of true, useful 2 × 300 MiSeq reads (derived from ~3 million raw reads) per sample, or ~0.75 million of true 2 × 100 HiSeq reads (derived from ~1.25 millions of raw reads) per sample. However, MiSeq datasets reach the plateau in ~1 815 mutations (Figure S1, yellow lines), while HiSeq saturates at ~580 to ~800 (Figure S1, blue lines). Only the sequencing platform can be invoked as source of differences since the same DNA was sequenced in both cases and the same bioinformatic pipeline run. It seems that MiSeq reactions would be more prone to sequencing variations than HiSeq reactions (in fact, we were verbally told by Illumina staff that the sequencing kit for 2 × 300 nt in MiSeq produces more sequencing errors than other kits). Therefore, being conservative, HiSeq could be considered true variations, while MiSeq is suspected to overestimate the genetic variation. The arrival of NovaSeq and the new sequencing platforms might benefit the performance of QuasiFlow. In any case, it seems reasonable to avoid sequencing mixtures from different platforms to measure genetic variability.

## DISCUSSION

The study of highly variable genetic systems was initiated in virology (Domingo and Schuster 2016) and now extends to other biological systems such as cancer (Dagogo-Jack and Shaw 2018), bacteria (Chan et al. 2013), mitochondria (Aryaman et al. 2019), or prions (Mahal et al. 2010). These analyses rely on accurate determination of mutations for the development of effective therapies. The aim of QuasiFlow (Fig. 1) is to provide sensitive, reliable, and robust analysis of complex genetic variability (including both low frequency mutations and recombinations) with a highly-stringent set of well-known bioinformatics tools glued with inhouse code. Results are easy to interpret when compared using QuasiComparer (Supplementary File 1) as illustrated with PCA and ANOVA plots grouping samples by their biological origin and not by the experimental process used to produce them (Figs. 2E-F, 3F and 4B-D). The capabilities of QuasiFlow/QuasiComparer couple have been demonstrated using synthetic sets of data (Fig. 2) and applied to a real-world data from plant viral (Fig. 3) and human mitochondrial (Fig. 4) populations.

The advent of the NGS (next-generation sequencing) technology has dramatically boosted the intrahost viral diversity studies, even if there is still no consensus about the most appropriate technology, the most appropriate software or the minimal read quality for SNV/recombination validations (McCrone and Lauring 2016). QuasiFlow applied to NGS data enabled, using a coverage >10 000× when a threshold of 20× was widely used in literature, the detection of low frequency SNVs and recombinations with a small rate of false positive detection. Emerging viruses such as coronavirus and geminiviruses are threatening humans and crops, respectively, due to their high frequency of mutation and recombination (Martin et al. 2011). Although some studies show that recombination is not the main phenomenon involved in the evolution of these viruses (Duffy et al. 2008), it is important to take this process into account due to its potential to generate large evolutionary jumps in very small time frames. Thus, recombinants can lead to host range expansion and increases in patogenicity. The analysis of the TYLCV-Mld infective clone in tomato plants in Fig. 3A-B demonstrated a striking triple infection (Fig. 3C), revealing that studies in greater depth of this type of co-infections could shed light on the reasons for fitness differences between such similar virus species.

The reliable analyses of the genetic heterogeneity of human mtDNA (Fig. 4) confirms the wide scope of QuasiFlow. In first instance, QuasiFlow confirmed the extreme genetic variation within mtDNA (Stewart and Chinnery 2021) with specific variant patterns in different tissues or nuclear backgrounds (i.e. sex), becoming a highly accurate and sensitive tool for the analysis of mitochondrial diseases. Interestingly, low-frequency mtDNA heteroplasmies were detected and then related to diseases that might have been elusive to detection by other means. Moreover, one can also envisage the utility of QuasiFlow/QuasiComparer in single-cell studies (Huang et al. 2021). In the case of tumours where genetic diversity is high from cell to cell due to a high mutation rate and such tumors can be considered as a collection of subclones, the study of this diversity can be facilitated using QuasiFlow: genetic diversity between patients or within a tumor can lead to pinpoint actionable driver mutations to design targeted therapies (Mroz and Rocco 2017; Baslan et al. 2020).

Accurately identifying mutations and variants among the thousands of virus that arise during an emerging viral infection can help understand how changes in tropism and jumps between host species occur. This is of particular importance in the present COVID-19 pandemic as fast haplotype reconstruction achieved by QuasiFlow can be a useful tool to detect variants of unknown origin such as Omicron that may be present as minority variants in mutant spectra.

## METHODS

### Workflow availability and requirements

QuasiFlow (Fig. 1), a workflow developed with the workflow manager AutoFlow (Seoane et al. 2016), takes as input raw, paired-end read data generated by a NGS platform (mainly Illumina, although other are allowed). Its design took advantage of computational resources available and the following dependencies: Art (Huang et al. 2012), SeqTrimBB, FastQC (Andrews 2010), BWA (Li and Durbin 2009), samtools (Li et al. 2009), Samstat (Li et al. 2009), Qualimap (Okonechnikov et al. 2015), ViReMa (Routh and Johnson 2014), VirVarSeq (Verbist et al. 2015) CliqueSNV (Knyazev et al. 2021), Varscan2 (Koboldt et al. 2012), VCFtools (Danecek et al. 2011), Haploclique (Töpfer et al. 2014), Clustal Ω (Sievers and Higgins 2014), CanvasXpress (https://github.com/neuhausi/canvasXpress/) and FactoMineR (Husson et al. 2009). It was coded and executed in Picasso supercomputer (3456 cores and 8.4 TB of RAM with Slurm queue system on SUSE Linux Enterprise Server 11SP2) at the University of Málaga. It requires (1) a reference genome in FASTA format (the begomovirus TYLCV or human mitochondrial DNA in this study), (2) a GFF file containing all open reading frame (ORF) coordinates, and (3) the circular or linear nature of the reference genome. Several samples can be run simultaneously using a customizable daemon script provided with the code that enables (1) executing QuasiFlow with all samples, (2) monitorizing QuasiFlow executions, (3) gathering results from all samples, and (4) comparative sample analysis using QuasiComparer. The workflow procceeds through the following seven tasks (Fig. 1):

#### Task 1: Read quality control and pre-processing

Raw reads were quality controlled using FastQC, and pre-processed using SeqTrimBB to remove low quality bases, vectors and adapters. SeqTrimBB default parameters were used, except for the following: 1) minimum phred base mean quality was set to 26 to avoid sequencing errors to be confounded with low frequency mutations and reduce the false positive ratio on ulterior analyses (Morelli et al. 2013; Welkers et al. 2014); 2) the minimum read length was increased to 100 nt to remove shorter reads confounding haplotype reconstruction (Zagordi et al. 2011). Viral reads were extracted from host and contaminant reads using the reference genome. This more reliable but reduced number reads (Fig 1, T1_U_rd) guaranteed soundness and robustness in further steps (Daly et al. 2015).

#### Task 2: Recombination events

The useful reads (T1_U_rd) were then treated as single reads (even though they come from paired-end reads) with ViReMa (Routh and Johnson 2014) using default parameters, excepting the aligner, which was configured to BWA (Li and Durbin 2009) instead of the default Bowtie since BWA is widely used for recombination studies. To avoid the identification of indels or misalignments as recombination events due to confounding reads, the seed minimum length was set to 55, the allowed mismatches was set to 1, and the microindels size was set to 55. Spurious recombination events with a frequency lower than 1 × 10^-3^ were removed to increase reliability in predicted recombination events. Additionally, recombination events located between the start and end positions of a circular reference sequence were discarded. Recombination events are classified by ViReMa as HomoRecs (Fig 1, T2_sR) for those resulting in a sequence deletion, or NonHomoRec (Fig 1, T2_dR) when the resulting sequence presents a duplication.

#### Task 3: SNV calling per ORF

The SNV calling process per ORF was based on VirVarSeq (Verbist et al. 2015) using useful reads in T1_U_rd and the GFF file provided at launch containing the ORF coordinates. VirVarSeq default parameters were used, but the minimum average phred quality is consistently set to 26 to assure that high quality reads are used. Variant codons were then converted to nucleotide positions (Fig1, T3_vcl) by assigning the SNV to the three nucleotides belonging to each codon.

#### Task 4: Mapping

Useful reads T1_U_rd (Fig. 1) were mapped on the genome reference to prepare data for subsequent haplotype reconstruction (Task 5) and variant calling (Task 6). To ensure high quality alignments, accurate mapping scoring (preventing false variant calling), and compatibility with further tools, mapping using BWA was launched with the following command:

~~~
bwa mem −B 20 −A 30 −O 30 −E 3 reference_sequence pair1.fastq pair2.fastq
| samtools view −bS −q 30 −F 4 > file.bam
~~~

A quality control analysis of the mapping is then performed on the BAM file with both Samstat (Li et al. 2009) and Qualimap (Okonechnikov et al. 2015) tools. Finally, by means of mpileup (Li et al. 2009), a first variant dataset is provided using the following command:

~~~
samtools mpileup −BQ 26 −q 30 −d 10000000 −f reference_sequence file.bam
~~~

where the −BQ flag filters by base quality (set to 26 in order to be coherent with the previous tasks). The −q flag, used in the current and in the previous samtools command, skips the alignments with a map quality of 30 or less. Finally, the −d flag changes the default maximum read depth from 8000 to 10^7^ following the samtools documentation ([CSL STYLE ERROR: reference with no printed form.]). Then, the pileup for one BAM file is saved (Fig 1, T4_pile).

#### Task 5: Haplotype reconstruction

The haplotype reconstruction relied on the recent CliqueSNV (Knyazev et al. 2021) that seems to outperform other haplotyping methods, even with low coverage samples where SNVs are at low frequencies. CliqueSNV opened each BAM file in T4_pile (Fig. 1) to identify groups of linked SNVs and remove sequencing errors. It resulted in a graph where SNVs are nodes and edges connect linked SNVs. Merging cliques in that graph identified haplotypes that were then saved (Fig 1, T5_Ht) together with their frequency.

#### Task 6: SNV calling

SNV calling was based on VarScan2 (Verbist et al. 2015). Mutations were called using the pileup file from Task 4 (T4_pile) with the following command

~~~
varscan mpileup2snp file.mpileup --min-coverage 10000 --min-reads2 10 --
min-var-freq 7e-6 --min-avg-qual 30
~~~

Indels were obtained with the same command but substituting mpileup2snp for mpileup2indel. The min-avg-qual removed low quality alignments, ensuring high quality calling SNVs, and the min-coverage and min-reads parameters selected only SNVs supported by 10 reads with a minimum read depth of 10 000. The min-var-freq parameter allows detection of real SNVs with frequencies as low as 7 × 10^-6^. The resulting files of mutations (Fig 1, T6_mut) and indels (Fig 1, T6_ind) were processed with the command vcf-stats from the VCFtools suite (Danecek et al. 2011) to obtain parameters such as variant counts, transition/transversion frequencies, as well as computing the frequencies of each type of base substitution that will be used below in QuasiComparer.

#### Task 7: Cross-validation and scoring

Taking in mind that haplotype reconstruction was poor when sequence divergence is low, failing to recover rare haplotypes (Schirmer et al. 2014), QuasiFlow was designed with a final task to remove unreliable haplotypes and increase the consistency with previously detected SNVs. To do so, haplotypes in T5_Ht were aligned to the genome reference using Clustal Ω, and the variable positions were cross-validated with previously obtained SNVs (T6_mut and T6_ind). Variability not supported by a SNV was edited, and haplotypes whose variations were all edited were then rejected as due to sequencing errors or untrusted variations. The resulting SNV-supported haplotypes (Fig. 1, T7_Supp_Ht) were realigned again and haplotype frequency was used to score diversity indices described by Gregori et al (Gregori et al. 2016), such as normalized Shannon, Simpson, Gini-Simpson, Hill numbers, sample nucleotide diversity, etc (Fig. 1, T7_DivI; see Supplementary Table S5 for a detailed description). In addition, haplotypes and SNVs were represented as a network where each haplotype was connected with all its SNVs to facilitate the interpretation of the genetic complexity and heterogeneity of the sample (see reports in Supplementary File 1). To facilitate result interpretation and visual user inspection an HTML report based on the javascript scientific library CanvasXpress (https://github.com/neuhausi/canvasXpress/) including the values for the parameters of Supplementary Table S5 as well as interactive plots and graphs is provided. It should be noted that most diversity indices (but Shannon) are provided for illustrative purposes since they are depending on the selected haplotypes.

### QuasiComparer

The variability analysis usually implies comparison of several samples or conditions. Since QuasiFlow can only manage one sequencing library (sample) at a time, inter-sample comparisons required the development of a dedicated comparative system called QuasiComparer. This analysis can be accomplished automatically by the same daemon script mentioned above provided that a description file with the following data is supplied: 1) the sample name, 2) a replicate identifier that allows to group the replicates that belongs to the same sample, and 3) the experimental group to which the sample is assigned, depending on the user experimental design (e.g., an experimental factor such as days post infection with three sample replicates taken at 7 days and other three replicates at 28 days).

QuasiComparer uses the parameters indicated in Supplementary Table S5 to perform correlation, ANOVA and principal component analysis (PCA) (Seoane et al. 2018) (Fig. 2B-G) to provide a new, interactive, compartive report in HTML (HTML reports included in Supplementary File 1) incluiding linear correlation plots, impact of each parameter on the experimental factors defined by the user by SNVs and by recombination events. The ANOVA is presented as a table and some circular plots (see Fig. 2F-G, one by each experimental factor described by the user) to illustrate the weight of each significant variable on each sample. The PCA also includes tables with information about its first two dimensions displaying the weight of each variable in PCA dimensions as well as plots (Fig. 4C), a PDF containing correlation analysis between the variables, and hierarchical clustering (Fig. 2E and 4D). All this information is intended to help users to see which samples have a different behavior in comparison to the rest, and correlate this behavior with the experimental factors used in the study.

### Synthetic and experimental datasets for testing

To test QuasiFlow and QuasiComparer performance, twelve independent synthetic read datasets were generated using Art (Huang et al. 2012) with the virus reference sequence of tomato yellow leaf curl virus Mild (TYLCV-Mld, AC# AF071228). The daemon was loaded with two parametres: 1) the presence of mutations and 2) the presence of recombinations, and called an in-house script that generates a FASTA file with mutated, recombinant and original virus sequences. Recombinations were generated making 30 recombinant events in a single sequence of TYLCV-Mld separated by 60 nt. The process was repeated until the recombinant sequences represented 20% of the total. To generate mutations, random changes were included in 10% of the nucleotide pool of TYLCV-Mld sequences. The FASTA file generated was then provided to Art, setting the read coverage to 300 000×, the maximum read length in 250 nt and a fragment size of 300 nt, with a standard deviation of 10 nt and using the MiSeq version 3.0 built-in error profile.

Four (mut_rec, mut_REC, MUT_rec and MUT_REC) synthetic sample datasets were generated by triplicate. “mut_rec” corresponded to reads where no mutation and no recombination were introduced and was considered as control group. “mut_REC” cotains reads where only recombinations were introduced and were considered as control for recombination. “MUT_rec” contains reads where only mutations were introduced, which makes it suitable as a control for mutation. Finally, the dataset “MUT_REC” contains reads where both mutations and recombinations were introduced and was considered the testing group.

As real-world dataset, an infectious clone of TYLCV-Mld was sequenced as a control before its passage through a plant to test the ability to detect mutations or recombination events introduced during high-throughput sequencing. This dataset was sequenced with a paired-end layout 2 × 300 nt resulting in 1 106 300 reads. After pre-processing and filtering to keep only TYLCV-Mld useful reads, this number decreased to 218 884.

### Samples, virus source and viral inoculation

Thirty day-old *Arabidopsis thaliana* Col-0 plants were inoculated with *Agrobacterium tumefaciens* LBA4404 containing the infectious clone (p1.7SP72/9) of TYLCV-Mld isolate [ES:72:97] (AC# AF071228). The construction harbours 1.7 genome-length of TYLCV-Mild (genome region from positions 161 to 1026 nt is only once) flanked by two intergenic (IR) regions that allow the excision of the viral monomer. Each *A. thaliana* plant was inoculated by the stem-puncture method using 20 μl of *A. tumefaciens* liquid culture adjusted to an OD600 of 1.00. After agroinoculation, plants were maintained for 28 days in a growth chamber (22 °C during the day and 20 °C at night, 70% relative humidity, with an 8-16 h day-night photoperiod of 8-16 h).

Dr. Enrique Moriones (IHSM “La Mayora”) kindly provided the tomato (*Solanum lycopersicum*) plants agroinoculated with TYLCV-Mld with disease symptoms and that were positive for the infection of begomoviruses of the TYLC-disease complex.

### DNA extraction, virus amplification, cloning, and sequencing

DNA from *A. thaliana* Col-0 and *S. lycopersicum* plants were extracted following the Edwards’ procedure (Edwards et al. 1991) with some modifications. From each sample, 150 mg of fresh tissue were ground with liquid nitrogen, and 400 μl of extraction buffer [200 mM Tris-HCl, 200 mM NaCl, 25 mM EDTA, SDS 0.5 % (w/v), pH 7.5, 0.1 % β-mercaptoethanol] added. After vortexing at 37 °C for 2 min, extracts were centrifuged at 12 000 × *g* at room temperature for 5 min. The supernatants were transferred to a new tube and mixed with 1 volume of phenol:chloroform (1:1). Following centrifugation at 12 000 × *g* at room temperature for 5 min, the aqueous phase was transferred to a new tube and mixed with 1 volume of isopropanol. After 2 min at room temperature, nucleic acids were recovered by centrifugation at 12 000 × *g* at room temperature for 5 min. The pellet was washed with 70% ethanol, dried and resuspended in 200 μl of RTE buffer (50 mM Tris-HCl pH 8.0, 10 mM EDTA pH 8.0, 0.1 % RNase A) to a concentration of 130 ng/μl by incubation at 65 °C for 10 min.

DNA samples from tomato and *A. thaliana* agroinfected plants were treated with DpnI endonuclease to remove *A. tumefaciens* viral DNA remnants but not viral DNA replicated in the plants. Endonuclease-treated DNA was purified with StrataClean Resin kit (Agilent Technologies), following the instructions given by the manufacturer.

Verification of the existence of the bioinformatically detected geminivirus species of tomato and agroinoculated *A. thaliana* plants was done by rolling-circle amplification (RCA) and restriction analysis. For this, 130 ng of DNA from young non-inoculated leaves were amplified for 18 h with the Illustra TempliPhi kit (GE Healthcare, UK) following manufacturer’s instructions. Amplified DNA was digested with single cutter (in viral genome) restriction enzyme BamHI to obtain the full-length DNA (2.7 kb). Restriction enzymes KpnI and ClaI were used to distinguish between TYLCV-Mld, TYLCV-Almería (AJ489258.1) and TYLCMaV (AF271234.1) in DNA extracts from tomato plants.

The presence of recombinant genomes between TYLCV-Almería and TYLCMaV in extracts form tomato plants was verified by PCR using reverse primer MAV-1271_1289-RW (5’-TTTTCTCAACTTCCGCATC) that binds to TYLCMaV sequence and forward primer IL-41_55-FW (5’-CCCACGAGGGTTCC) that binds to TYLCV-Almería sequence. RDP4 program (Martin et al. 2015) was used to analyse the recombination event of the most frequent haplotype. Digestions and PCR results were analysed by agarose (Agarose SPI, Duchefa) gel electrophoresis 1 % (w/v) with TBE buffer.

To generate DNA for Illumina sequencing viral DNA from total DNA plants extracts and the infectious clone of TYLCV-Mld [ES:72:97] were amplified by RCA. To avoid creating a population bottleneck during the RCA amplification step two microliters of the plant DNA extract (260 ng total) were used as template. Replicates of each amplified sample were pooled to reduce the accumulation of random errors due to Phi29 Polymerase. Phi29 polymerase has an associated 3’-5’ exonuclease “proofreading” activity and has a reported error rate of 3-5 × 10^-6^. The contribution of the Phi29 polymerase error during RCA is about 100-fold lower than the observed mutation frequency of begomoviruses (around 10^-4^ mutations per nucleotide or higher).

MiSeq and HiSeq sequencing was performed at Macrogen (Seoul, South Korea). For MiSeq, equimolar sequencing libraries of tomato and nine *A. thaliana* samples were done with Truseq DNA PCR-free (550 bp insert) kit and sequenced on an Illumina MiSeq as 2 × 300 bp reads. For HiSeq, libraries were done for three *A. thaliana* samples with Truseq DNA PCR-free (350 bp insert) kit and sequenced on an Illumina HiSeq2000 as 2 × 100 bp reads. The number of reads belonging to geminiviruses that exceeded the initial quality filter was 293 668, with an average coverage of 14 772× per nucleotide.

### Mitocondrial sequences for genetic variability analyses

Human mtDNA sequences BioProject PRJEB5005 contained blood, lymphoblastoid cell lines and saliva samples from males and females. The set comprised 47 libraries of 2 × 300 bp paired-end reads obtained with the Illumina MiSeq platform with a number of reads between 1.38 × 10^5^ and 4.32 × 10^5^. Samples from the same tissue and sex were considered biological replicates. The Revised Cambridge Reference Sequence (“rCRS”) of 16 569 nucleotides (AC# NC_012920) was used as standard reference for human mtDNA. Since the mean coverage was about 1500×, variants with a frequency below 9 × 10^-4^ (0.09 %) were filtered out by means of QuasiFlow parameters --min-var-freq to 9×10e-4 and the --min-coverage to 1000 in Task 6. The biological significance of mitochondrial variants was obtained using MITOMAP database (Lott et al. 2013) (https://www.mitomap.org). To assess tissue origin differences between the lymphoblastoid cell lines and blood a *t*-Student was performed using FactoMineR for each axis of the PCA.

## DATA ACCESS

### Data

Sequencing reads can be found at BioProject PRJNA823046. Human mtDNA sequences were obtained from BioProject PRJEB5005.

### Code

QuasiFlow, QuasiComparer and the daemon script (daemon.sh) that controls their execution can be downloaded from https://bitbucket.org/seoanezonjic/quasiflow/src/master/. The resulting reports are attached as Supplementary File 1.

## ACKNOWLEDGEMENTS

This work was supported by Consejería de Economía, Innovación y Ciencia, Junta de Andalucía with assistance from the European Regional Development Fund (ERDF) and the European Social Fund (ESF) [grant to Research Groups BIO-264 and BIO-267; P10-CVI-6561 and UMA18-FEDERJA178 to A.G.-P., E.V.M; UMA20-FEDERJA-029 to MGC; and a predoctoral fellowship to L.D.-M. A.G-P. acknowledges support of Plan Propio de Investigación y Transferencia of the University of Málaga. Open access charges was funded by University of Málaga. The authors also thankfully acknowledge the computer resources and the technical support provided by the Plataforma Andaluza de Bioinformática of the University of Málaga.

## AUTHOR CONTRIBUTIONS

PS designed QuasiFlow and QuasiComparer and constructed the synthetic data. LDM analysed the TYCLV and mitochondrial datasets. EV interpreted the mitochondrial data. MGC supervised the algorithm implementation and execution, and wrote the manuscript. AGP conceived the research, supervised the viral interpretations, and wrote the manuscript.

## SUPPLEMENTARY INFORMATION

**Supplementary File 1** (DOI <u>10.6084/m9.figshare. 19518976</u>): Compressed zip file containing the following HTML reports

- *1-mut_rec.html:* QuasiFlow report for mut_rec synthetic reads.
- *2-MUT_REC.html:* QuasiFlow report for MUT_REC synthetic reads.
- *3-Comparative_Report_SyntheticReads.html:* QuasiComparer comparative report for all synthetic reads
- *4-Control_TYLCV.html:* QuasiComparer comparative report for the control of TYCLV-Mld replicated in vitro.
- *5-Tomato_TYLCV-invected.html:* QuasiFlow report for TYCLV samples from tomato-infected cells.
- *6-Comparative_Report_Mitochondria.html:* QuasiComparer comparative report for mtDNAs.

**Supplementary File 2** (DOI <u>10.6084/m9.figshare.19519003</u>): Complementary data related to:

- *Table S1:* Evaluation of TYLCV-Mld genetic variation.
- *Table S2:* Nucleotide changes detected in the mtDNA samples used to validate QuasiFlow analyses.
- *Table S3:* mtDNA variants detected that have a known relation with a human disease.
- *Table S4:* ANOVA test for the 47 samples considering the different factors evaluating genetic variation provided by QuasiFlow
- *Figure S1:* Plot of the number of SNPs detected depending on the number of nucleotides analysed. Nine MiSeq datasets (yellow lines; ranging from 115 to 335 million reads) and three HiSeq dataset (blue lines, ranging from 500 to 501.5 million reads) were used. Every dataset was randomly subset in bulks of 50 000 read folds and plotted against the number of SNPs detected in it by Varscan2.
- *Table S5:* Variables calculated during QuasiFlow execution

